# A synthesis of over 9,000 mass spectrometry experiments reveals the core set of human protein complexes

**DOI:** 10.1101/092361

**Authors:** Kevin Drew, Chanjae Lee, Ryan L. Huizar, Fan Tu, Blake Borgeson, Claire D. McWhite, Yun Ma, John B. Wallingford, Edward M. Marcotte

**Affiliations:** Center for Systems and Synthetic Biology, Institute for Cellular and Molecular Biology, University of Texas at Austin, Austin, TX 78712, USA; Department of Molecular Biosciences, University of Texas at Austin, Austin, TX 78712, USA; The Otolaryngology Hospital, The First Affiliated Hospital of Sun Yat-sen University, Sun Yat-sen University, Guangzhou, China

**Author notes:** Current address: Recursion Pharmaceuticals, Inc., Salt Lake City, UT 84108, USA.

**Keywords:** Human interactome, mass spectrometry, proteomics, protein complexes, cilia, ciliopathy

## Abstract

Macromolecular protein complexes carry out many of the essential functions of cells, and many genetic diseases arise from disrupting the functions of such complexes. Currently there is great interest in defining the complete set of human protein complexes, but recent published maps lack comprehensive coverage. Here, through the synthesis of over 9,000 published mass spectrometry experiments, we present hu.MAP, the most comprehensive and accurate human protein complex map to date, containing >4,600 total complexes, >7,700 proteins and >56,000 unique interactions, including thousands of confident protein interactions not identified by the original publications. hu.MAP accurately recapitulates known complexes withheld from the learning procedure, which was optimized with the aid of a new quantitative metric (*k*-cliques) for comparing sets of sets. The vast majority of complexes in our map are significantly enriched with literature annotations and the map overall shows improved coverage of many disease-associated proteins, as we describe in detail for ciliopathies. Using hu.MAP, we predicted and experimentally validated candidate ciliopathy disease genes *in vivo* in a model vertebrate, discovering CCDC138, WDR90, and KIAA1328 to be new cilia basal body/centriolar satellite proteins, and identifying ANKRD55 as a novel member of the intraflagellar transport machinery. By offering significant improvements to the accuracy and coverage of human protein complexes, hu.MAP (http://proteincomplexes.org) serves as a valuable resource for better understanding the core cellular functions of human proteins and helping to determine mechanistic foundations of human disease.

## Introduction

A fundamental aim of molecular biology is to understand the relationship between genotype and phenotype of cellular organisms. One major strategy to understand this relationship is to study the physical interactions of the proteins responsible for carrying out the core functions of cells, since interacting proteins tend to be linked to similar phenotypes and genetic diseases. Accurate maps of protein complexes are thus critical to understanding many human diseases (Goh et al. 2007; P. I. Wang and Marcotte 2010; Lage et al. 2007). Technical advances in the field of proteomics, including large-scale human yeast two-hybrid assays (Rolland et al. 2014; Rual et al. 2005), affinity purification / mass spectrometry (AP-MS) (Huttlin et al. 2015; Hein et al. 2015) and co-fractionation / mass spectrometry (CF-MS) (Havugimana et al. 2012; Wan et al. 2015; Kirkwood et al. 2013; Kristensen, Gsponer, and Foster 2012) have enabled the partial reconstruction of protein interaction networks in humans and other animals, markedly increasing the coverage of protein-protein interactions across the human proteome. Such efforts are largely ongoing, as we still lack a comprehensive map of human complexes, and we have only partial understanding of the composition, formation and function for the majority of known complexes. Prior high-throughput protein interaction assays in yeast and humans have generally tended to show limited overlap (Yu et al. 2008; von Mering et al. 2002; Gandhi et al. 2006; Hart, Ramani, and Marcotte 2006), suggesting that interactions from different studies tend to be incomplete, possibly error-prone, but also orthogonal.

Over the past year, three large-scale protein interaction mapping efforts in particular have greatly expanded the set of known human protein interactions, namely BioPlex (Huttlin et al. 2015), Hein *et al*. (Hein et al. 2015) and Wan *et al*. (Wan et al. 2015), collectively comprising 9,063 mass spectrometry shotgun proteomics experiments. The three resulting datasets are notable for representing independent surveys of human protein complexes by distinct methods (AP-MS vs. CF-MS), in distinct samples (different cells and tissues), and in the case of the two AP-MS datasets, using distinct choices of affinity-tagged bait proteins. The datasets are complementary in other aspects as well: The two AP-MS interaction sets are each sampled from a single choice of immortalized cancer cell line grown in rich cell culture medium, and thus represent deep, but condition-and cell type-specific, views of the interactome network. The AP-MS networks sample only a fraction of human proteins as “baits” and are limited to interactions which contain a bait protein, which is expressed recombinantly as a fusion to an affinity purification moiety (green fluorescent proteins (GFP) for Hein *et al.* or FLAG-HA for BioPlex.) These strategies resulted in 23,744 and 26,642 protein interactions for BioPlex and Hein *et al*., respectively. In contrast, the CF-MS experiments sampled endogenous proteins in their native state without genetic manipulation, but with only partial purification, relying instead on repeat observation of co-eluting proteins across samples and separations to increase confidence in the interactions. The resulting 16,655 protein interactions reflect the biases expected for well-observed proteins, tending towards more abundant, soluble proteins. Additionally, the Wan *et al*. interactome required all interactions to have evidence in at least two sampled metazoan species; thus, only evolutionarily-conserved human proteins are represented. As a consequence, none of these three datasets is individually comprehensive; nonetheless, we expect them to present highly complementary, potentially overlapping views of the network of core human protein complexes. There is thus an opportunity to integrate these over 9,000 published mass spectrometry experiments in order to create a single, more comprehensive map of human protein complexes.

Here, we describe our construction of a more accurate and comprehensive global map of human protein complexes by re-analyzing these three large-scale human protein complex mass spectrometry experimental datasets. We built a protein complex discovery pipeline based on supervised and unsupervised machine learning techniques that first generates an integrated protein interaction network using features from all three input datasets and then employs a sophisticated clustering procedure which optimizes clustering parameters relative to a training set of literature curated protein complexes. While generating the complex map, we re-analyzed AP-MS datasets to identify >15,000 high-confidence protein interactions not reported in the original networks. This re-analysis substantially increased the overlap of protein interactions across the datasets and revealed entire complexes not identified by the original analyses. Importantly, the integrated protein interaction network and resulting complexes outperform published networks and complex maps on multiple measures of performance and coverage, and represents the most comprehensive human protein complex map currently available. Moreover, the framework we employ can readily incorporate future protein interaction datasets.

We expect that a comprehensive definition of protein complexes will ultimately aid our understanding of disease relations among proteins. In line with expectation, our map shows markedly increased coverage of disease-linked proteins, especially for proteins linked to ciliopathies, a broad spectrum of human diseases characterized by cystic kidneys, obesity, blindness, intellectual disability, and structural birth defects (Hildebrandt, Benzing, and Katsanis 2011). We highlight both known and novel complexes relevant to ciliopathies and, moreover, experimentally validate multiple new protein subunits of ciliary complexes, using *in vivo* assays of cilia structure and function in vertebrate embryos. Additionally, we distribute our results to the community in a simple and easy to navigate website: http://proteincomplexes.org/. The scale and accuracy of this human protein complex map thus provides avenues for greater understanding of protein function and better disease characterization.

## Results and Discussion

### Overlap between three recent high-throughput animal protein interaction datasets is modest, but can be greatly increased by a re-analysis of the data

Protein interaction networks from various sources often show minimal overlap (von Mering et al. 2002; Gandhi et al. 2006; Hart, Ramani, and Marcotte 2006). We therefore first sought to measure the overlap of proteins and interactions between three recently published protein interaction data sets from BioPlex (Huttlin et al. 2015), Hein and colleagues (Hein et al. 2015) and Wan and colleagues (Wan et al. 2015). The BioPlex network is the result of 2,594 AP-MS experiments from HEK293T cells. Similarly, the Hein *et al.* network is the result of 1,125 AP-MS experiments from HeLa cells. In both screens, the authors considered only interactions between the affinity-tagged bait protein and the co-precipitated “prey” proteins, corresponding to a “spoke” model of interactions (Figure 1A). The Wan *et al.* network is derived from a CF-MS analysis of nine organisms, comprising 6,387 MS experiments.

**Figure 1:**
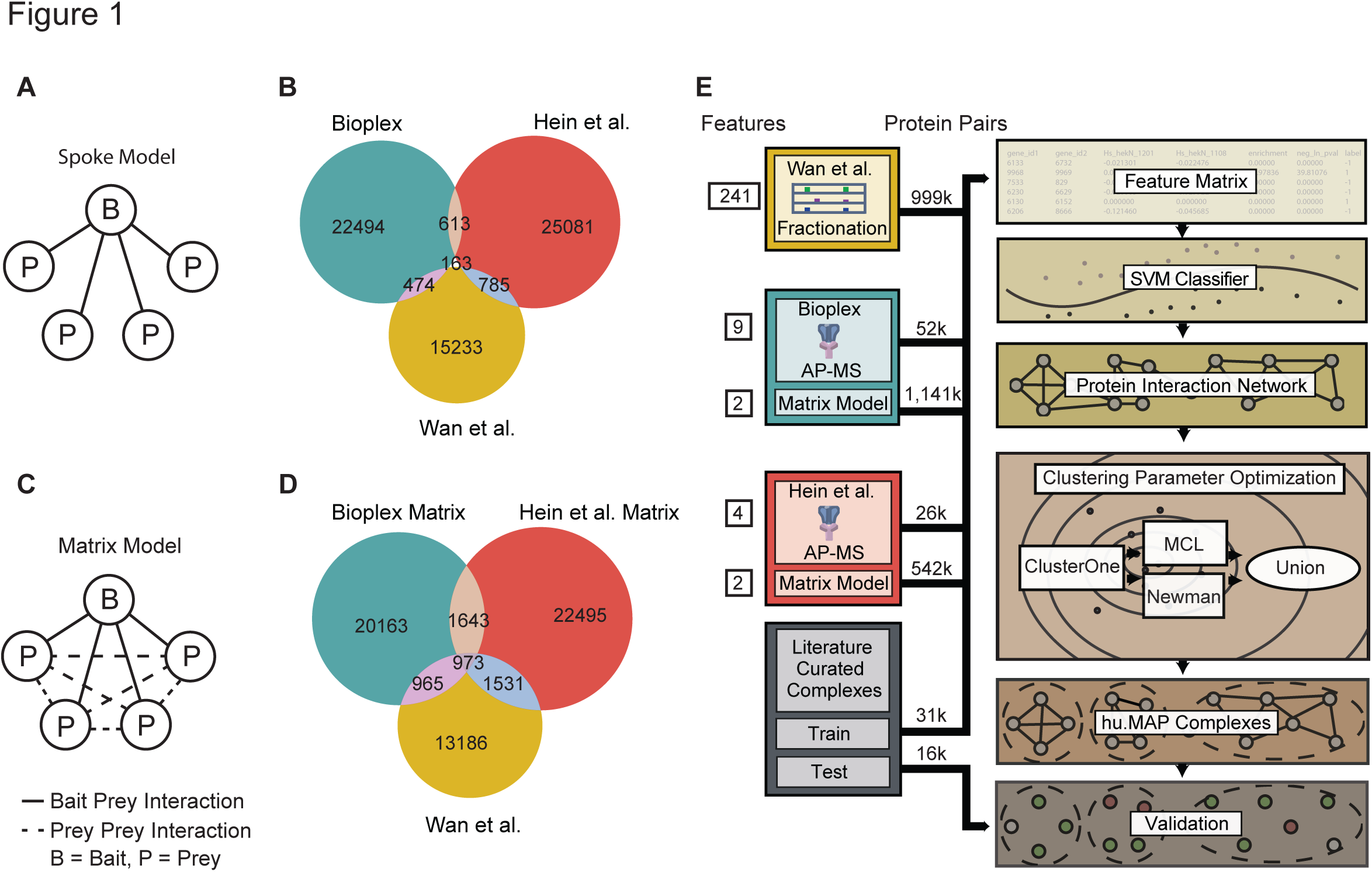
Re-analysis of published AP-MS experiments improves overlap among protein interaction networks. A. Graphical schematic of spoke model applied to AP-MS datasets. In the spoke model, all interactions must include a bait protein. B. Venn diagram of overlap between published large scale protein interaction networks BioPlex (AP-MS), Hein *et al*. (AP-MS) and Wan *et al*. (CF-MS). Protein interactions in BioPlex and Hein *et al*. were generated from a spoke model. C. Graphical schematic of matrix model applied to AP-MS datasets. In the matrix model interactions are allowed between prey proteins. D. Venn diagram of overlap between protein interaction networks where a matrix model was applied to BioPlex and Hein *et al*. Sizes of matrix model protein interaction networks were kept constant with published networks. Note an increase in the overall number of overlapping interactions when compared to B. E. Diagram of protein complex discovery workflow. Three protein interaction networks, BioPlex, Hein *et al*. and Wan *et al.* were combined into an integrated protein complex network and clustered to identify protein complexes. Parameters for the SVM and clustering algorithms were optimized on a training set of literature curated complexes and validated on a test set of complexes.

We observe reasonable overlap in terms of the proteins identified within each published network, ranging between 30% and 68% of the proteins between individual networks (**Supplemental Table 1**). However, the overlap among protein interactions was more limited, ranging between ~3% and ~6% overlap (Figure 1B, Supplemental Table 1). There are generally three accepted reasons for the limited overlap commonly observed between large-scale protein interaction maps (von Mering et al. 2002): 1) the interaction networks sample different portions of the interactome (e.g., differences in cell types and baits), 2) the experimental methods used are biased towards discovery of certain classes of interactions (e.g. soluble vs. membrane protein interactions) and therefore are complementary to the other methods, and 3) the experimental methods produce false positive interactions.

To further probe the reason for the limited observed overlap, we next considered if the spoke model interpretation of the AP-MS experiments was partly responsible. By only considering interactions between bait proteins and their preys, spoke models are heavily reliant on the baits selected for experimentation, and also ignore evidence for repeated precipitation of intact complexes across baits. Traditionally, spoke models have shown higher accuracy when compared to the alternative full “matrix” model interpretation (Figure 1C) (Bader and Hogue 2002). However, the discrimination between true and false protein interactions can be dramatically improved by computing confidence scores for prey-prey interactions when applying a matrix model (Hart, Lee, and Marcotte 2007; H. Wang et al. 2009) or a hybrid spoke-matrix model (e.g., socio-affinity index) (Gavin et al. 2006) to AP-MS data. In order to reinterpret the AP-MS datasets using a matrix model while effectively discriminating true and false positive interactions, as well as suppressing “frequent flyer” co-purifying proteins, we applied a hypergeometric distribution-based error model to the AP-MS datasets, calculating *p*-values for pairs of proteins that were significantly co-precipitated more often than random across AP-MS experiments. We then ranked each protein pair according to its calculated *p*-value and selected the top N pairs for each AP-MS dataset, where N is the number of interactions reported in the original published interaction networks (23,744 BioPlex interactions and 26,642 Hein *et al.* interactions). This reinterpretation of AP-MS experiments using a matrix model substantially increased the amount of overlap among the three interaction networks, which rose to between 10% and 15%, as plotted in Figure 1D (see also **Supplemental Table 1**). This result indicates that there are thousands of interactions captured by the AP-MS experiments that were not previously identified and confirms a far greater consistency among the underlying mass spectrometry datasets, arguing that a combined analysis of the datasets could considerably improve coverage of the complete human protein interactome.

### Integrating the large-scale proteomics datasets into a human protein-protein interaction network

Based on the notion that considering this large and diverse set of experiments jointly should increase the ability to discriminate between true and false protein interactions, we next asked if integrating all three large-scale datasets would outperform the individual networks in terms of identifying true human protein interactions. We employed a formal machine-learning framework to combine evidence from the thousands of individual mass spectrometry experiments in the three large-scale datasets. Our approach was specifically designed to address the limited network overlap described above, using the confidence-weighted matrix model to increase interaction coverage while preserving accuracy. We expected the orthogonal techniques employed, CF-MS and AP-MS, to complement each other, where CF-MS captures stable interactions among endogenous proteins in diverse cells and tissues, while AP-MS captures a large collection of interactions with differing biophysical characteristics. The three datasets also sample very different portions of the human interactome in terms of cell type and bait selection, which we similarly expected to contribute to a more comprehensive map.

Figure 1E outlines the pipeline used for protein complex discovery. We first generated a feature matrix using the published features from BioPlex, Hein *et al.* and Wan *et al.* as well as the new matrix model features, in the form of a negative log hypergeometric *p*-value capturing the specificity and extent to which pairs of proteins co-precipitated across many AP-MS baits. Rows in the feature matrix represented pairs of proteins and columns represented measured numerical estimates of protein pairs’ interaction potentials based on the different experiments (see Materials and Methods). We also labeled protein pairs according to their support by a gold-standard, literature-curated set of human protein complexes (the CORUM protein complex database (Ruepp et al. 2010)). We assigned a positive label if both proteins were seen in the same complex, a negative label if both proteins were observed in the literature-curated set but not in the same complex, and an “unknown” label for all other pairs. A support vector machine (SVM) classifier was trained using the labeled feature matrix, then applied to all protein pairs, assigning each pair an SVM confidence score, indicating the level of support for that pair of proteins to participate in the same complex. This classification step thus resulted in an integrated human protein-protein interaction network, in which the nodes are proteins identified in any of the three experimental datasets, and the edges between nodes represent co-complex interactions weighted proportionally to the SVM score.

As an initial estimate of the quality of the integrated human protein interactions, we calculated their precision and recall by reconstructing a set of 15,687 gold-standard, literature-curated co-complex interactions omitted from the training procedure. While networks generated using features from only one of the three datasets showed high precision for high confident interactions, they quickly dropped in precision in the higher recall range (Figure 2A). In contrast, the integrated network demonstrated substantial improvements to performance, with a precision of 80% over just under half of the benchmark interactions. Additionally, adding the matrix model features to the published interactions greatly improved the performance, indicating that the matrix model features capture new information beyond spoke features and serve as a rich source of evidence supporting true protein interactions.

**Figure 2:**
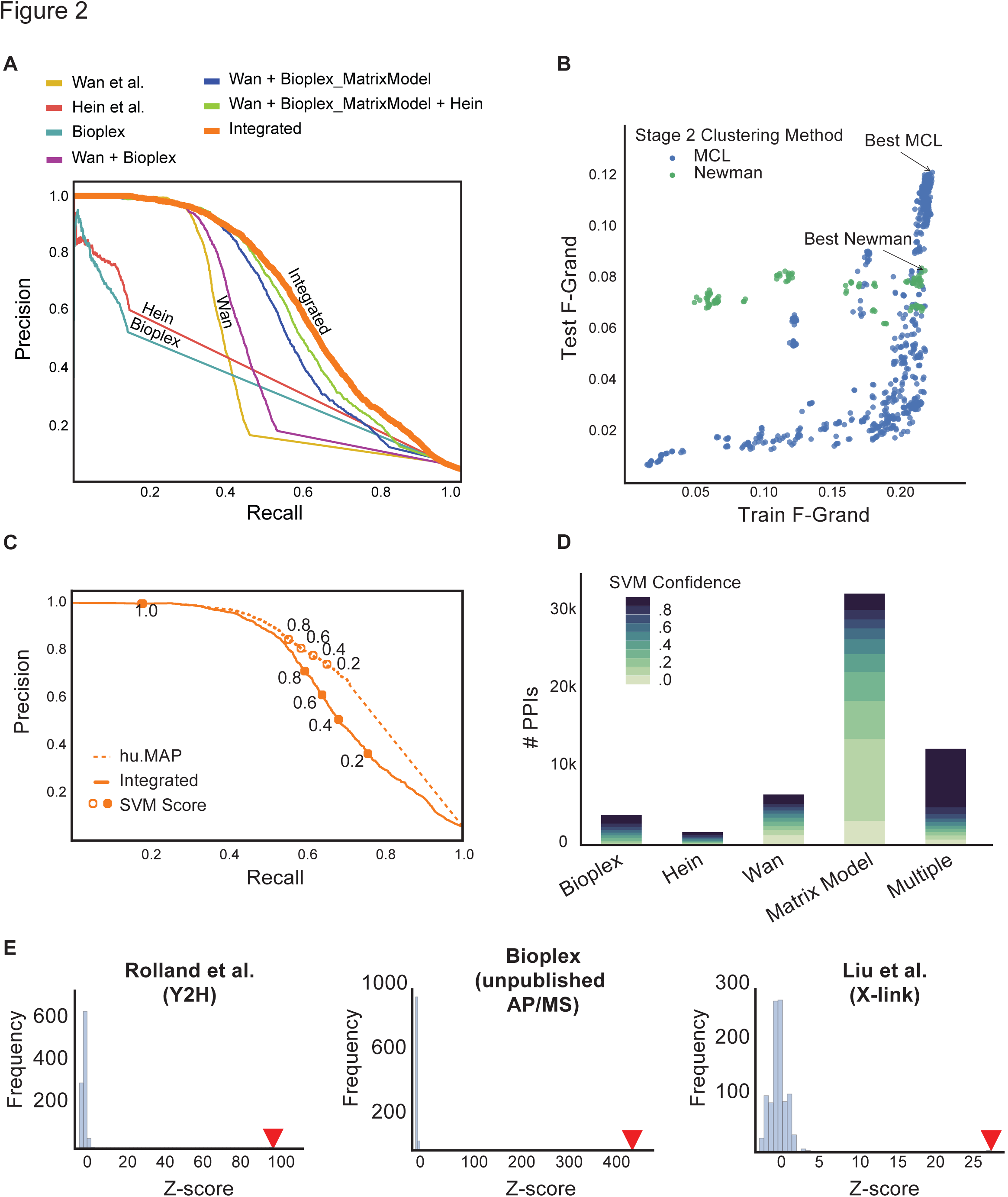
Integration of the three large-scale protein complex datasets substantially improves both precision and recall of known human protein interactions. A. Precision-recall curves calculated on a leave-out set of protein interactions from literature curated complexes for different combinations of predictive protein interaction features. The integration of all three datasets outperforms all other networks. Also, note a substantial improvement in performance when the matrix model features are used (Wan+BioPlex vs. Wan+BioPlex_MatrixModel). B. Performance of parameter optimization for MCL and Newman two-stage clustering procedures. Each data point represents a set of parameters and is evaluated based on the resulting clusters similarity to both training and test sets of complexes using the F-Grand measure (see Materials and Methods). Final parameter sets were selected based only on F-Grand measure for the training set. C. Precision-recall curves evaluating protein interactions on leave out set before (integrated) and after (hu.MAP) clustering procedure. Note an improvement in performance after clustering suggests the clustering procedure successfully removed false positive interactions. D. Distribution of protein interactions in the final protein interaction network based on input evidence. Note the matrix model interactions produce many high confident interactions. Also, the “Multiple” category shows predominately high confident interactions, which validates the integration of multiple datasets mitigating false positives. E. Protein interactions from our complex map significantly overlap with other protein interaction datasets across a variety of experimental types.

### Clustering pairwise interactions reveals human protein complexes

A hallmark of protein complexes is that their component proteins should frequently be co purified in independent separations and affinity purifications. This trend manifests as densely connected regions of the interaction network, which we sought to identify by applying a two stage clustering procedure. In the first stage of clustering, we applied the ClusterOne algorithm (Nepusz, Yu, and Paccanaro 2012), which identifies large, dense subnetworks of the full protein interaction network. Importantly, ClusterOne allows proteins to participate in more than one sub network as dictated by the data, as proteins frequently participate in more than one complex (Wan et al. 2015). In the second stage, we separately applied MCL (Enright, Van Dongen, and Ouzounis 2002) and Newman’s hierarchical clustering method (Newman 2004) to further refine the sub-networks produced by ClusterOne. As with many unsupervised machine learning techniques, clustering algorithms have adjustable parameters for optimizing their performance. We therefore used a parameter sweep strategy to identify choices of parameters that best recapitulated known complexes. We evaluated each parameter combination by comparing the resulting protein clusters to our literature curated training set of protein complexes and selected the top ranking parameter combination. As the comparison of protein complexes to a gold standard set is not a fully solved problem, we first developed an objective scoring framework for complex-level precision and accuracy, as we describe in the next section.

### Evaluation of the derived protein complexes using k-cliques identifies the most accurate and comprehensive complex map

Guiding and assessing the accuracy of the reconstructed complexes requires comparison with a gold standard set of known complexes. However, comparing sets of complexes to known complexes (or more generally, comparing sets of sets with each other), is ill-defined due to the problem of first deciding which sets should be compared and second, to the incomparable nature of specific matches. For instance, given two non-overlapping complexes, one of size 3 and one of size 20, it is difficult to assess whether an exact match of the complex of size 3 should be given more weight than a partial match of the complex of size 20 (e.g., with 17 out of 20 correct). Many complex-complex comparison metrics (Song and Singh 2009; Brohée and van Helden 2006; Bader and Hogue 2003) have attempted to address this issue, but they are often difficult to interpret and may lead to false minima in the parameter landscape, in part because they require a mapping procedure to determine which specific gold standard complexes match up with specific reconstructed complexes.

In order to more systematically address these issues, we invented a new class of similarity metrics, *k-cliques*, for comparing sets of complexes in a formal precision-recall framework. Specifically, our approach is based on the matching of cliques within the set of all possible cliques between predicted complexes and benchmark complexes. Cliques range from size 2 (pairwise protein interactions) through *n*, where *n* is the size of the largest predicted complex. The approach allows for precision and recall values to be calculated unambiguously (because a clique is either present or absent in a given set) for each clique size, *k*, and averaged to determine a single performance metric (here, the F-Grand metric, corresponding to the average across all clique sizes of the harmonic mean of precision and recall; see Materials and Methods). An important feature of the *k*-cliques approach is that it focuses the evaluation on protein interactions, rather than the proteins themselves, providing a unique perspective on set comparisons. In addition, while other approaches suffer from evaluating each complex individually and often require a cluster reduction step in which similar clusters are combined to avoid potential skew, e.g. as caused by prediction of sub-complexes of larger complexes, the *k*-cliques approach compares complexes on a global level and naturally deals with potential skew by only evaluating on the unique set of cliques for all predicted complexes. Finally, there is no need to determine a unique mapping between each predicted and benchmark complex, thus avoiding mapping-induced ambiguity.

We computed the performance in terms of reconstructing known complexes for each of >1,000 different clustering algorithm parameter combinations, varying the SVM confidence threshold for the input pairwise protein interactions, the ClusterOne density and overlap options, and the inflation option for MCL. The top-scoring sets of clusters for the two 2^nd^ stage clustering methods, MCL and Newman’s hierarchical method, were of similarly high quality when evaluated relative to the training set of complexes (Figure 2B). These two top-scoring cluster sets also showed the top-ranking scores when compared to the literature-curated leave-out test set for their respective clustering methods, serving to validate the parameter optimization method. As the two top-scoring cluster sets identified many distinct specific complexes and sub-complexes, we combined these two top scoring definitions of complexes in order to provide a more comprehensive view of the myriad of physical protein assemblies in human cells. The resulting fully integrated human protein complex map, called hu.MAP, consists of 4,659 complexes, 56,735 unique co-complex interactions and 7,777 unique proteins (**Supplemental Tables 2 and 3**).

### The integrated map improves pairwise interaction performance, identifies new interactions, and is strongly supported by independent protein interaction datasets

We wished to assess the quality of the integrated map of human protein complexes by multiple, independent approaches. First, because the process of network clustering entails removing interactions between proteins that are inconsistent with the defined complexes, we might expect the resulting clustered network to be more accurate than the pre-clustered network. Indeed, the final interaction network shows improved precision and recall (Figure 2C), indicating that the clustering step is preferentially removing false positives from the original network.

Next, during the course of identifying protein interactions and complexes, we withheld a leave out set of literature-curated complexes to serve as a final, fully independent test set. We compared these data to the derived map and to previously published complex maps, using two different comparison measures (Supplemental Figure 1). For both the *k*-clique metric and the precision recall product measure (Song and Singh 2009), we observed a dramatic improvement in performance over the Wan *et al.* and BioPlex maps (note: Hein reported only interactions, not complexes). A survey of evidence supporting each interaction in the map showed multiple lives of evidence supported many pairwise interactions (Figure 2D). This further supports the notion that the underlying datasets are orthogonal and that integrating them provides substantial improvement on discriminating true and false protein interactions. Remarkably, however, we observed tens of thousands of interactions in the map supported only by matrix model features, 15,454 of them having very high confidence. Thus, considering prey-prey interactions in the AP-MS datasets dramatically enhanced the identification of human protein interactions.

Finally, in order to assess the quality of the final map independently of both the test and training set complexes, we further evaluated our complex map with several of the largest remaining available human protein interaction datasets. We observed highly significant overlap with protein interactions from different experimental methods, including yeast two-hybrid assays (Rolland et al. 2014), additional unpublished BioPlex AP-MS experiments (“BioPlex” 2016), and crosslinking mass spectrometry performed on human cell lysate (Liu et al. 2015) (Figure 2E). Thus, comparisons with independent datasets strongly support the high quality of the derived protein complexes, as measured by multiple metrics of performance, considering interactions both pair-wise and set-wise, and even considering interactions measured independently by multiple different technologies.

### Prey-prey interactions reveal a large, synaptic bouton complex, isolated from HEK cells

The thousands of additional high-confidence interactions contributed by prey-prey co-purification patterns led us next to consider their value in our protein complex discovery pipeline. In particular, we asked if matrix model edges could independently identify complexes, or if they only served to support observed bait-prey associations. We thus searched for complexes in the map that were supported predominantly with matrix model interactions. Figure 3A summarizes AP-MS experiments for four example complexes. Three of these complexes—the exosome complex, eukaryotic initiation factor 3 (eIF3) complex, and the 19S proteasome—were supported both by spoke edges and matrix model edges, showing high complementarity between the two sets of interactions. This support was evident in the strong interaction density both between bait proteins and between bait and prey proteins within each complex. In contrast, the fourth complex shown in Figure 3A is a newly identified complex by our pipeline that surprisingly has limited density between bait proteins, but substantial, high-specificity density in the prey region of the matrix. Notably, the four bait proteins that each precipitate nearly all 60 subunits of this complex largely do not co-precipitate each other.

**Figure 3:**
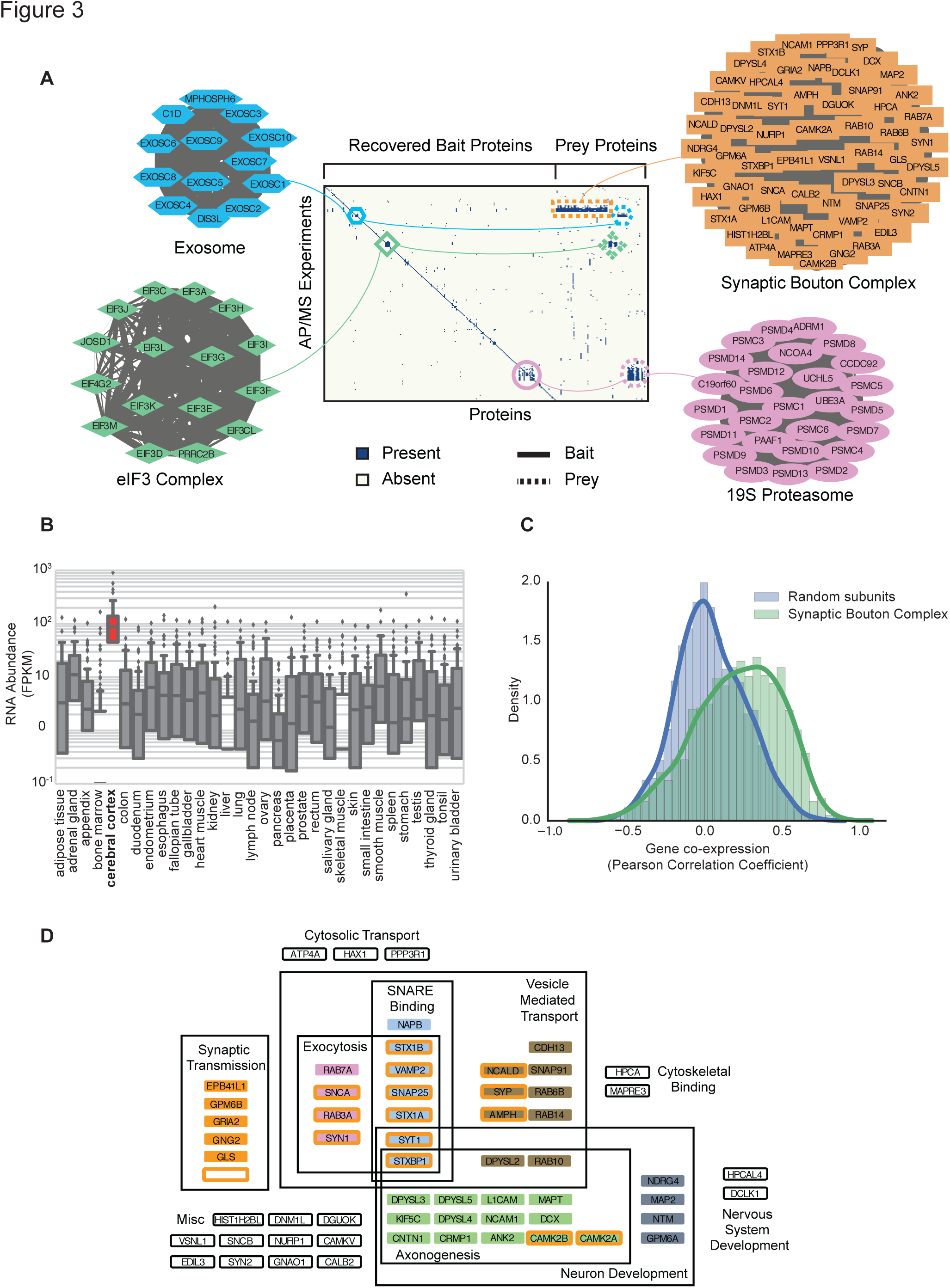
Matrix model edges identify large synaptic bouton complex. A. Presence/absence matrix of BioPlex AP-MS experiments as rows and pulled down proteins as columns for four complexes identified in our complex map. The Exosome, eIF3 Complex and 19S Proteasome all have multiple bait-bait interactions whereas the novel synaptic bouton complex does not have bait-bait interactions but does have substantial density in the non-bait region of the matrix. This density is identified by the matrix model and highlights the model’s ability to discover protein complexes. B. RNA expression profiles of proteins in the synaptic bouton complex across different tissues sampled by the Human Protein Atlas. This shows the complex is highly specific for cerebral cortex tissue. C. Correlation coefficient distributions of Allen Brain Map tissue expression profiles between synaptic bouton complex proteins and random set of proteins. This shows coherence of expression among proteins in the complex suggesting a functional module. D. Significantly enriched Gene Ontology annotations for proteins in the synaptic bouton complex shows enrichment for neuron development and synaptic transmission.

We performed annotation enrichment analysis to establish functional connections between member proteins of this novel complex. Strikingly, the proteins identified in this complex are highly specific for cerebral cortex tissue, as measured by Human Protein Atlas tissue expression data (Uhlén et al. 2015) (Figure 3B). We additionally observed high brain-region specific co-expression among members of the complex, unlike as for random protein pairs, in the Allen Brain Map microarray dataset (Hawrylycz et al. 2012) (Figure 3C). The complex includes subunits of the SNARE complex, a known physically associated set of proteins involved in synaptic vesicles (Südhof 1995). Consistent with this trend, we found a strong enrichment of gene ontology terms (GO) (Ashburner et al. 2000) among members of the complex specific to neurotransmission and neuron migration (Figure 3D). Thus, there is good correspondence between this complex and known interacting protein complexes at the synaptic bouton, the presynaptic axon terminal region containing synaptic vesicles and the location of neuronal connections.

Rather surprisingly, the AP-MS experiments that support this complex were all performed with HEK293T cells. HEK293T cells were first reported to be derived from human embryonic kidney tissue (Graham et al. 1977), and therefore it is puzzling as to why a complex comprised of cerebral cortex specific proteins showed such a strong signal in kidney-derived cells. However, re-analyses of HEK293T cell origins suggest that they were originally mis-annotated and actually derive from adjacent human embryonic adrenal tissue, rather than embryonic kidney cells (Shaw et al. 2002; Lin et al. 2014), and thus exhibit many neuronal properties (Shaw et al. 2002). The possibility remains open that the protein complex identified here could also have additional roles in the body. Regardless, this complex exemplifies the value of prey-prey interactions for discovering protein complexes.

### The integrated map markedly improves coverage of disease-linked protein complexes

A key application of more accurate human protein complex maps will be to highlight and characterize biologically important protein modules, especially those relevant to human disease. We thus next evaluated the map in reference to a variety of localization, functional and disease annotation datasets. First, we annotated proteins in hu.MAP with information about their human tissue expression patterns from the Human Protein Atlas (Uhlén et al. 2015). We observed a substantial portion of proteins in our map expressed across all assayed tissues, suggesting our map captures many core processes in human cells (Figure 4A), although many tissue-specific complexes appear to be identified as well, as for the example of the synaptic bouton complex in Figure 3. We next evaluated the fraction of complexes with significantly enriched annotations (Bonferroni-corrected hypergeometric *p* < 0.05; g:Profiler (Reimand et al. 2016)) from the Gene Ontology, Reactome, CORUM, OMIM, KEGG and HPA annotation databases (Ashburner et al. 2000; Fabregat et al. 2016; Ruepp et al. 2010; Amberger et al. 2015; Kanehisa et al. 2014; Uhlén et al. 2015) (Figure 4B). Approximately two-thirds (3,147 out of 4,659) of the complexes had at least one significantly enriched annotation term, demonstrating the biological pertinence of complexes in the map (see **Supplemental Table 4** for full list of each complexes’ significantly enriched annotation terms).

**Figure 4:**
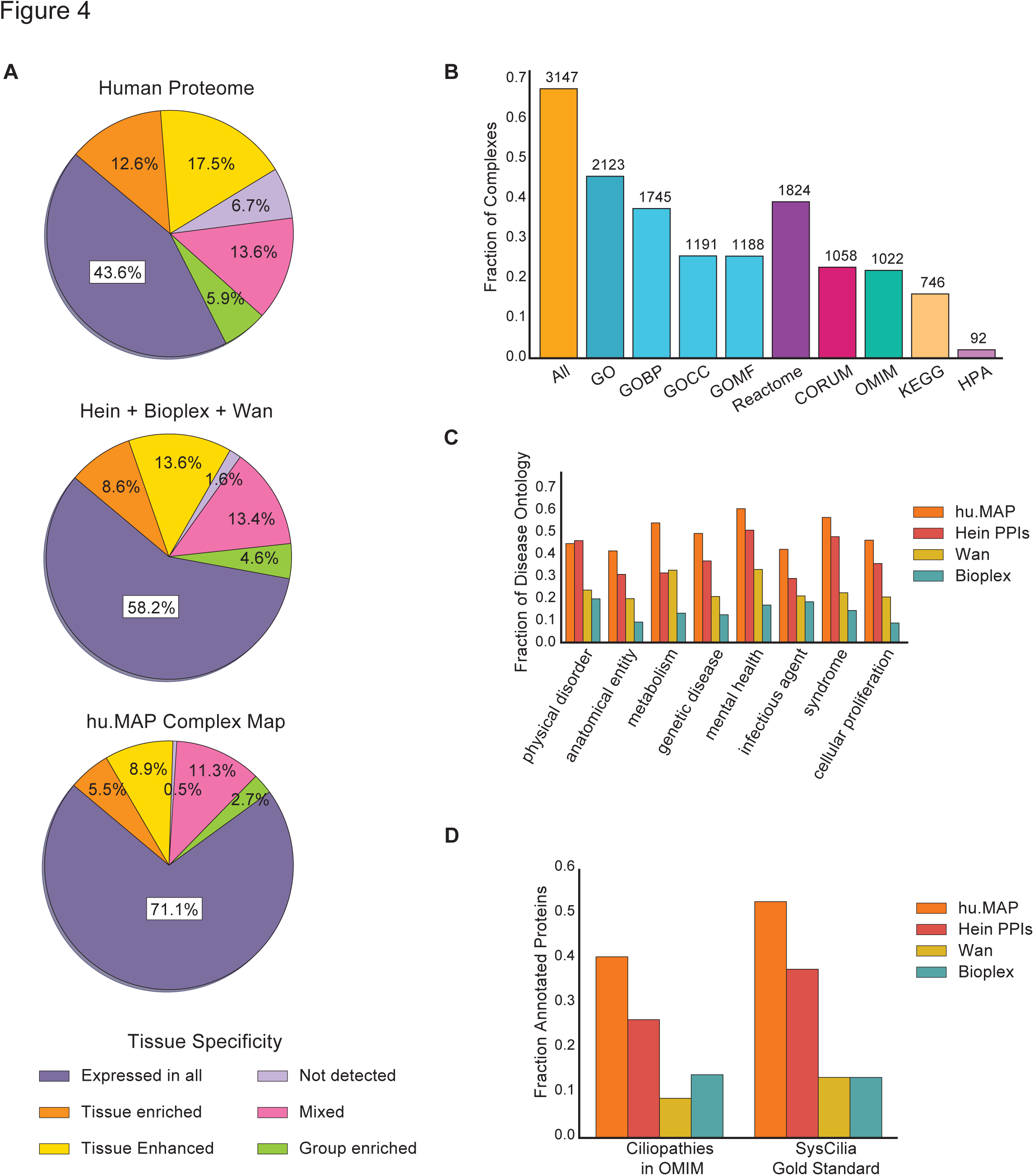
hu.MAP consists of predominately core human complexes and covers a large fractions of disease genes. A. Complex map coverage of Human Protein Atlas RNA tissue specificity classifications showing majority of complexes are ubiquitously expressed and likely core cellular machinery. B. Fraction of complexes with significantly enriched annotation terms (p-value < 0.05, g:Profiler hypergeometric test with Bonferroni correction) from various ontologies. C-D. Protein coverage of high-level Disease Ontology terms (C) and cilia related annotations (D) for complex map as well as two published maps (Wan *et al.* and BioPlex) and a published interaction network (Hein *et al.*).

Knowledge that a protein interacts with a disease-associated protein greatly increases the probability that the first protein is linked to the same disease (Dudley et al. 2005; Ideker and Sharan 2008; Fraser and Plotkin 2007; McGary, Lee, and Marcotte 2007; Lage et al. 2007). Thus, we expect an important application of this map will be to enable the discovery of candidate disease genes. In order to estimate this strategy’s potential, we compared the map’s coverage of known disease-associated proteins with other published networks. Figure 4C shows the fraction of proteins annotated in the Online Mendelian Inheritance in Man (OMIM) disease gene database, mapped according to eight high level Disease Ontology (DO) terms (Schriml et al. 2012) for hu.MAP, the Wan map, BioPlex, and the full Hein *et al.* interaction network (which serves to increase its proteome coverage). hu.MAP shows substantially higher coverage than the other networks for nearly all high level DO terms, covering ~46% of the annotated human disease-associated proteins.

### New components of ciliary protein complexes

One specific class of diseases in particular stood out, namely diseases related to defective cilia, known as ciliopathies. Cilia are microtubule based cellular protrusions that are critical for cell-to-cell signaling and proper embryonic development (Oh and Katsanis 2012; Goetz and Anderson 2010). Cilia assembly and maintenance are highly regulated processes whose disruption can lead to debilitating birth and early childhood disorders, including Joubert syndrome, Meckel syndrome, Bardet-Biedl syndrome, orofaciodigital syndrome, and polycystic kidney disease. Although many ciliopathies share clinical presentations such as kidney and liver dysfunction, other clinical features and their severity can vary considerably across individuals (Tobin and Beales 2009; Gerdes, Davis, and Katsanis 2009; Hildebrandt, Benzing, and Katsanis 2011). The resulting confounding array of clinical features, an absence of cures, and limited but expensive treatments, all lead ciliopathies to collectively represent a major health burden (Tobin and Beales 2009). Protein complexes are integral to many ciliary and centrosomal processes and have major roles in ciliopathies (Gupta et al. 2015; Boldt et al. 2016). To more directly assess hu.MAP’s relevance to ciliopathies, we measured its coverage of ciliopathy-associated proteins (OMIM-annotated proteins mapped onto the mid-level Disease Ontology term “ciliopathy”) and known ciliary proteins (literature-curated as the SysCilia “Gold Standard” (van Dam et al. 2013)) (Figure 4D). For both ciliopathy-associated and ciliary proteins, we observed a substantial increase in coverage over other networks, with hu.MAP covering >50% of ciliary proteins.

An examination of individual complexes enriched with ciliary proteins highlighted both known and novel ciliary components. hu.MAP reconstructed multiple known ciliary protein complexes including the Intraflagellar Transport particles A and B (IFT-A and IFT-B) (Cole et al. 1998; Piperno and Mead 1997), the Bardet-Biedl-linked BBSome (Nachury et al. 2007), the CPLANE ciliogenesis and planar polarity effector complex (Toriyama et al. 2016), and the CEP290-CP110 complex (Tsang et al. 2008) (Figure 5). In all, the map contains 234 complexes and sub-complexes involving 158 ciliary proteins (**Supplemental Table 5**), many associated with ciliopathies (Toriyama et al. 2016; Chetty-John et al. 2010; Walczak-Sztulpa et al. 2010; Schaefer et al. 2014; Beales et al. 2007; den Hollander et al. 2006). Moreover, we observed many of these complexes to also contain additional uncharacterized proteins. These novel proteins represent excellent candidates for ciliary roles, so we next focused on detailed experiments to characterize their *in vivo* functions and subcellular localization in developing vertebrate embryos.

**Figure 5:**
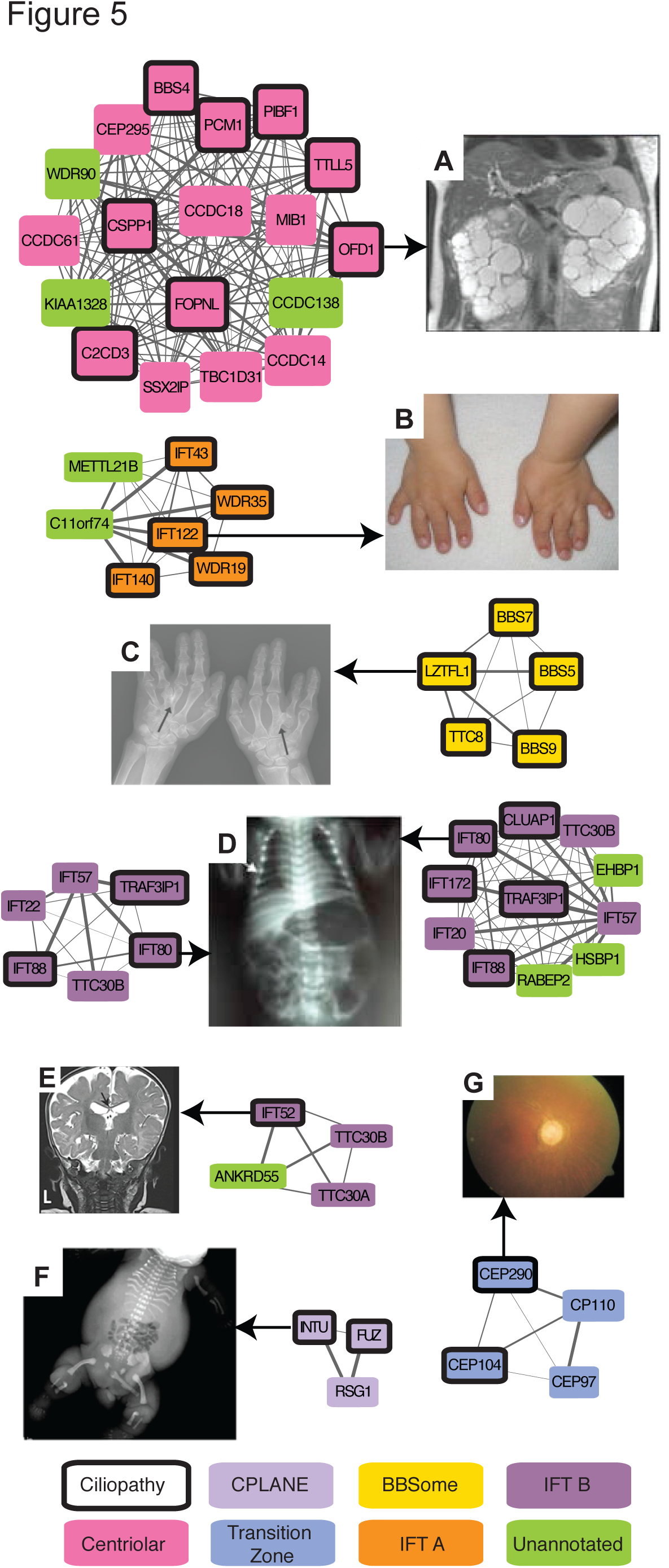
Select complexes in the map are strongly linked to human ciliopathies. Eight complexes are highlighted with ciliopathy-linked subunits (bold outlines), illustrating representative clinical presentation of associated mutations as follows: A. Polycystic kidneys from patient with OFD1 variant, adapted from (Chetty-John et al. 2010). B. Brachydactyly in Sensenbrenner syndrome patient with IFT122 variant, adapted from (Walczak-Sztulpa et al. 2010). C. X-ray of hands of Bardet–Biedl syndrome patient with LZTFL1 (BBS17) variant, adapted from (Schaefer et al. 2014). D. X-ray exhibiting chest narrowing of Jeune asphyxiating thoracic dystrophy individual with IFT80 variant, adapted from (Beales et al. 2007). E. Brain MRI showing hypoplastic corpus callosum from patient with an IFT52 variant, adapted from (Girisha et al. 2016), F. X-ray of short-rib polydactyly syndrome individual with multiple skeletal anomalies with INTU variant, adapted from (Toriyama et al. 2016). G. Undeveloped fovea among other pathologies in Leber congenital amaurosis patient with CEP290 variant (den Hollander et al. 2006).

### Observation of an 18 subunit ciliopathy-linked complex enriched in centrosomal proteins

Among the ciliary complexes, we identified a large, 18-subunit complex in which 8 subunits were already linked to ciliopathies and 14 members were known to localize to the centrosome centriolar satellites (Figure 5A and 6A). A second 8-member complex was observed interacting with subunits of the first complex, also containing centrosome-localized and ciliopathy-linked proteins (Figure 6A). Figure 6B plots the AP-MS observations that supported the discovery of these complexes. We observed strong evidence for physical associations among members in each complex, with multiple bait proteins from multiple datasets affinity-purifying substantial portions of each complex. Centrosomes are the microtubule organizing centers of cells, with dual roles in chromosomal movement and organization of the ciliary microtubule axonemes. Thus, the marked enrichment of centrosomal/centriolar satellite and ciliopathy proteins in these two complexes strongly implicates a relationship between centriolar satellites and ciliary related disease.

**Figure 6:**
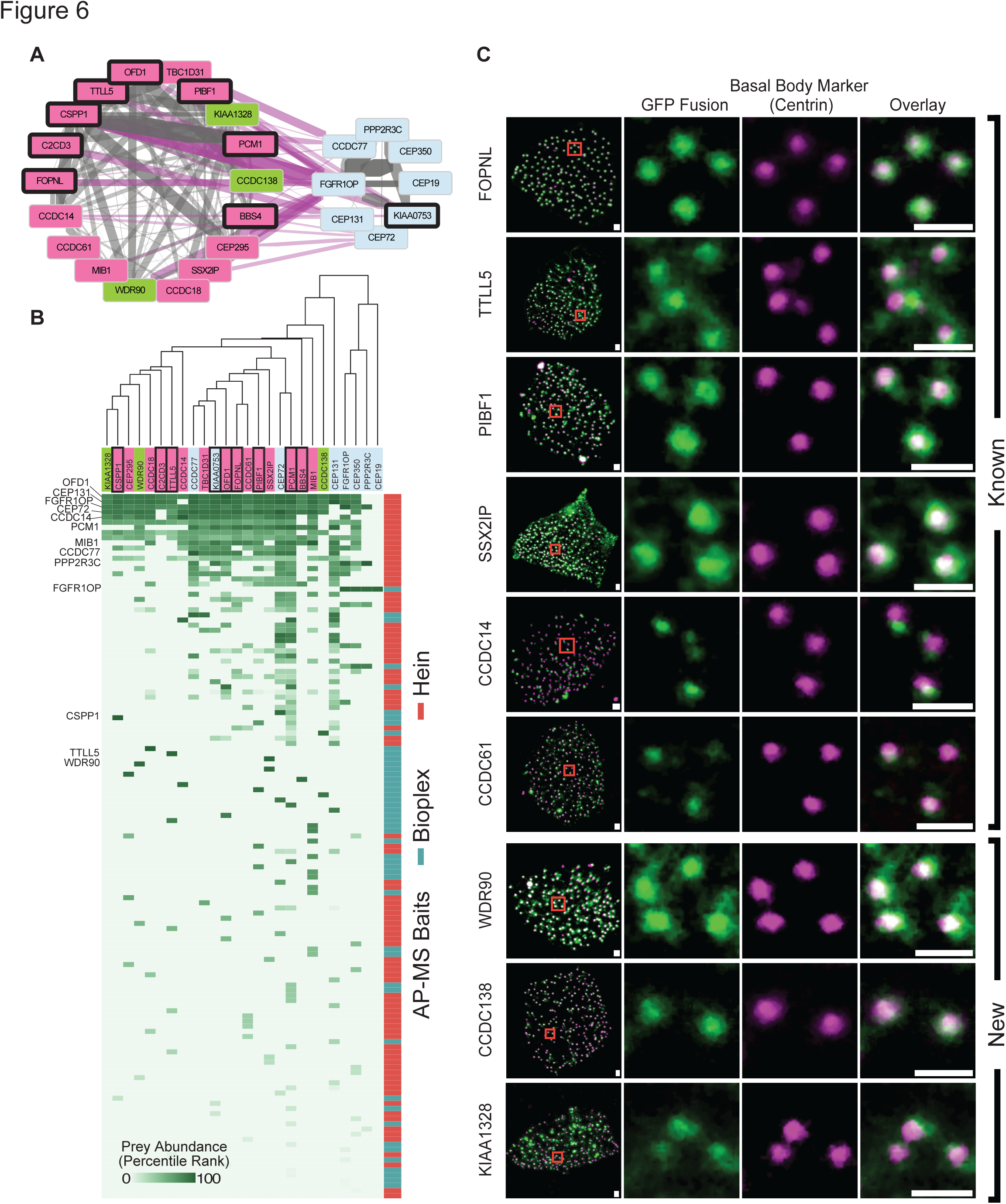
Oro-facial-digital syndrome 1 (OFD1) interaction partners are centriole and centriolar satellite proteins, suggesting new components of ciliary basal bodies. A. Network of Ciliopathy Complex and closely interacting centrosomal complex. Edge weights represent SVM confidence scores where gray are intra-complex edges and purple are inter-complex edges. Color of nodes follows **Figure 5** conventions. B. Matrix of AP-MS evidence supporting both complexes. The matrix shows strong support for interactions within each complex. Bait proteins that are members of either complex are labeled on the left. C. Experimental validation of ciliary proteins using multi-ciliated epithelial cells in *Xenopus laevis*. Localization assays for the three uncharacterized proteins in the OFD1 complex confirm that all three proteins localize to basal bodies at the base of the cilia in a manner similar to known components of the complex. Scale bars: 1um.

Three of the 18 proteins in the larger complex were completely uncharacterized (WDR90, CCDC138 and KIAA1328), so we determined the subcellular localization of tagged versions of these proteins as a direct experimental test of the map’s prediction. We expressed proteins in this complex as GFP-fusions in multiciliated cells (MCCs) of embryos of the frog *Xenopus laevis*, as these cells provide an exceptional platform for studying vertebrate ciliary cell biology *in vivo* (Werner and Mitchell 2012; Brooks and Wallingford 2012; Toriyama et al. 2016). Serving as positive controls, known centrosomal components, including PIBF1, localized strongly and specifically to basal bodies and co-localized with the basal body marker Centrin4 (Figure 6C). WDR90, CCDC138 and KIAA1328 each localized strongly and specifically to basal bodies, strongly supporting their participation in centrosomal and ciliary biology, and validating the map’s predictions.

### ANKRD55 is a novel intraflagellar transport complex protein

We next focused on the IFT complexes, which link cargos to microtubule motors for transport along ciliary axonemes (Taschner and Lorentzen 2016). The IFT system is comprised of two multi-protein complexes, IFT-A and IFT-B (Cole et al. 1998; Piperno and Mead 1997). Our map effectively recapitulated known protein-protein interactions in the IFT-B complex, assembling not only the entire complex (Figure 5D, Supplemental Figure 2), but also recovering elements of known sub-complexes. For example, the map assembled much of the known IFT-B “core” (also called the IFT-B1 complex) containing IFT22, IFT46, IFT74 and IFT81. The map also identified a complex containing IFT38, IFT54, IFT57, and IFT172, which closely matches the recently described IFT-B2 complex (Taschner et al. 2016; Katoh et al. 2016). The map further recapitulated the smaller IFT-A complex (Figure 5B, Supplemental Figure 2), the anterograde IFT motor complex of KIF3A, KIF3B and KAP (Taschner and Lorentzen 2016), and also more ancillary but relevant interactions, such as that between IFT46 and the small GTPase ARL13B (Cevik et al. 2013).

Importantly, the map also predicted novel components of the IFT complexes. For example, the map predicted an interaction between IFT-B and RABEP2 (Figure 5D), which is interesting because while RABEP2 is implicated in ciliogenesis (Airik et al. 2016), its mechanism of action remains obscure. Even more interesting is the link between IFT-B and the poorly defined protein ANKRD55 (Figures 5E and 7A). A re-examination of the raw data from the AP-MS experiments reinforced the notion that ANKRD55 is an IFT-B component (Figure 7B), so we tested this hypothesis *in vivo* using high-speed confocal imaging in *Xenopus* MCCs (Brooks and Wallingford 2012). We find that an ANKRD55-GFP fusion protein localizes to cilia and moreover time-lapse video analysis indicates that ANKRD55 traffics up and down cilia (Figure 7C and **Supplemental Movie 1**). In kymographs made from the time-lapse data, we observed ANKRD55-GFP to move coordinately in axonemes with known IFT protein CLUAP1-RFP (Supplemental Figure 3A and **Supplemental Movie 2 and 3**). Finally, disruption of the ciliopathy protein JBTS17 was recently shown to elicit accumulation of IFT-B proteins (but not IFT-A proteins) in ciliary axonemes (Toriyama et al. 2016). Consistent with the predicted association of ANKRD55 with IFT-B, we observed robust aberrant accumulation of ANKRD55 in axonemes after *JBTS17* knockdown (Supplemental Figure 3C).

**Figure 7:**
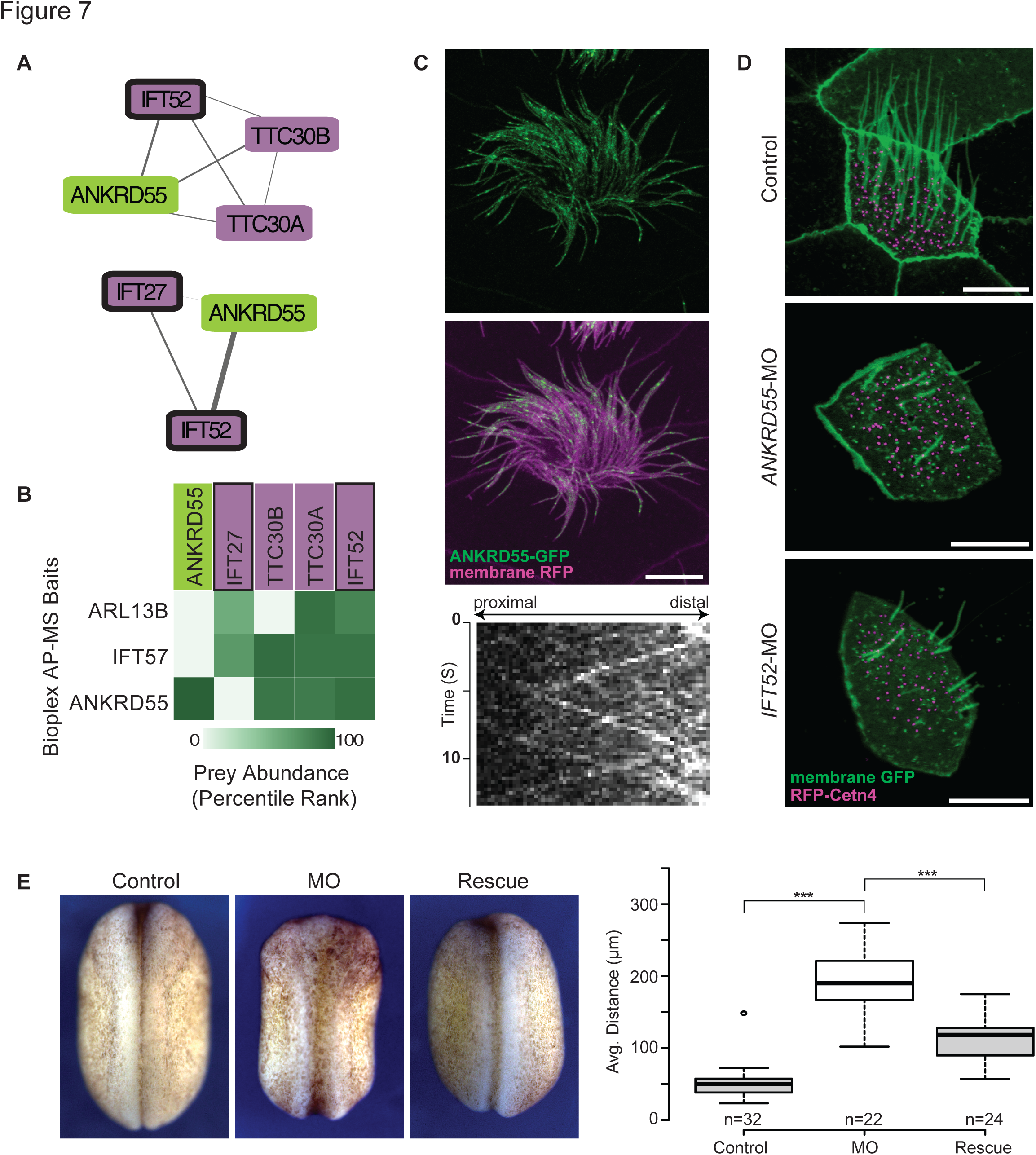
ANKRD55 is a new component of the intraflagellar transport (IFT) particle, is important for ciliogenesis, and has a role in neural tube closure. A. Network view of two IFT subcomplexes associated with ANKRD55. B. Matrix of AP-MS experiments shows strong support for ANKRD55 association with known IFT proteins. C. ANKRD55 localizes to cilia as predicted from co-complex interactions, as assayed *in vivo* in multi-ciliated *X. laevis* epithelial cells. Scale bar: 10 um. Kymograph of ANKRD55 localized to cilia *in vivo* reveals rapid trafficking along the length of the cilia. D. Morpholino knockdown of *ANKRD55* results in reduced count and length of cilia, in a manner similar to the control IFT52 knockdown, supporting a role in ciliogenesis for ANKRD55. Scale bar: 10 um. E. Dorsal view of stage 19 embryos displays that *ANKRD55* knockdown causes neural tube closure defects that are rescued by wild-type *ANKRD55* mRNA. The box plot displays average distance between neural folds in control, morphant, and rescue embryos. *** (P < 0.0001)

Because IFT subunits have been linked to vertebrate birth defects, as a new subunit of the IFT-B particle, we would expect disruption of ANKRD55 to in turn disrupt ciliary function and proper embryonic development. We performed *in vivo* experiments in order to test ANKRD55 function, first by asking if knockdown elicited a similar cilia phenotype to IFT knockdown. Figure 7D shows images of morpholino-antisense oligonucleotide (MO) knockdowns of both *ANKRD55* and its co-complex member *IFT52*; the morphant embryos exhibit similar ciliary disruption phenotypes, further supporting the connection between IFT and ANKRD55. Finally, disruption of IFT in vertebrate animals, including *Xenopus* and mice, result in defects in neural tube closure (Huangfu et al. 2003; Toriyama et al. 2016), and we therefore asked if loss of ANKRD55 would exhibit similar defects. Indeed, knockdown of *ANKRD55* in *Xenopus* embryos resulted in defective neural tube closure; this defect could be rescued by expression of a version of the *ANKRD55* mRNA that could not be targeted by the MO, arguing for the specificity of the knockdown phenotype (Figure 7E). Taken together, the interaction, localization, and genetic perturbation data all indicate that ANKRD55 interacts physically and functionally with the IFT-B complex and strongly suggests that ANKRD55 is likely to play a role in human ciliopathies.

## Conclusions

Gaining a more complete understanding of the relationship between human genotypes and phenotypes will require improved maps of protein complexes as well as some understanding of their dynamic nature across cell types and across the spectrum from healthy to diseased tissue. Recent advances in proteomics now allow for the comparison of biological networks across different conditions to identify the dynamics of protein complex function (Ideker and Krogan 2012; Kristensen, Gsponer, and Foster 2012). However our ability to interpret these experiments is hindered by the lack of a complete picture of protein complexes. Here, we report a map that captures a significant portion of the core protein machinery in human cells. This map provides not only a framework on which to organize future experiments, but also provides immediate insight into broad classes of human diseases, including ciliopathies.

To produce this map, we described the re-analysis and integration of three large-scale protein interaction datasets. We showed that the limited overlap of the input published networks is due in part to the computational analyses of the underlying experiments, which suggests more sophisticated analysis techniques may further uncover novel protein interactions. Integration across the datasets greatly enhanced the precision and recall of the final interaction network, in part by scoring prey-prey interactions, leading us to identify thousands of interactions which were previously unreported in the original publications, as for e.g. the synaptic bouton complex. This weighted matrix model approach should be of increasing importance because of its ability to elegantly compensate for and capitalize on off-target identifications in AP-MS datasets. The model’s ability to take into account the frequency at which proteins are identified across experiments allows for the filtering out of non-specific and contaminating proteins found across datasets. It is likely the matrix model approach will only become more powerful as additional datasets are available and can be combined to identify subtle trends across many experiments.

Indeed, there is tremendous effort in the community to generate ever larger-scale maps of human protein interactions, and extensions to ongoing high throughput interactome studies can be naturally incorporated into our protein complex discovery framework. We envision a continual expansion and refinement of this set of human protein complexes using the described pipeline as new high throughput protein interaction experiments are published.

We also developed a novel method for comparing the reconstructed protein complexes to a gold standard set of protein complexes, a problem that has proven difficult for the field. The solution we propose is formulated in a precision-recall framework based on cliques derived from the predicted clusters and gold standard set. This approach differs from previous solutions in that it generates a global comparison between clusters and the gold standard, rather than identifying the best match for each single cluster at a time. The method is applicable whenever one wishes to compare two sets of sets, as it is general in nature and should be useful beyond comparing protein complexes.

The success of long-standing efforts to understand the genetic basis of human disease relies heavily on understanding the physical interactions of proteins. We demonstrate the value of our complex map for understanding human disease by featuring ciliopathy related complexes. Through this analysis we highlighted uncharacterized proteins, which we experimentally validated to be cilia-associated, as predicted by the map. We also knocked down one of these proteins, ANKRD55, and showed a disruption in ciliogenesis, which strongly suggests a role in ciliopathies. These results establish the ability of an integrated human protein complex map to identify new candidate disease genes, with potentially broad applicability to many human diseases.

## Materials and Methods

### Gold standard training and test set complexes

For training and evaluating our protein complex discovery pipeline we used literature curated complexes from the Corum core set (Ruepp et al. 2010). We first removed redundancy from the Corum set by merging complexes that had large overlap (Jaccard coefficient > 0.6). The set of complexes were then randomly split into two sets, labeled test and training. Due to proteins participating in multiple complexes, the randomly split sets were not fully disjoint. We dealt with the overlap of these two sets differently at the pairwise interaction and complex level, as follows:

For the purposes of training and evaluating our SVM classifier, we generated positive and negative pairwise protein interactions for both test and training sets. A positive protein interaction is defined as a pair of proteins that are part of the same complex. A negative protein interaction is defined as a pair of proteins that are both in the set of complexes but not part of the same complex. We addressed overlap here between the test and training (positive/negative) protein interactions by removing interactions from the training protein interaction sets which were shared in the test protein interaction sets, such that the sets were fully disjoint.

For the analyses of protein complexes, in order to ensure that the test and training sets of complexes were disjoint, we removed entire complexes from the training set which shared any edge with a complex in the test set. In comparing the size distributions (the number of subunits per complex) between the training and test sets, we noticed a skew of larger complexes in the test set likely a result of our conservative approach of removing complexes from the training set. In order to better balance the training and test complex set size distributions, we first randomly split the test set into two and combined one half with the training complexes. We again applied our redundancy removal procedure, removing complexes from the training set which shared any edge with a complex in the test set. Similar to what has been done previously (Wan et al. 2015; Havugimana et al. 2012), we also removed complexes larger than 30 subunits from the test set so as not to skew performance measurements.

The final pairwise protein interaction training/test sets consisted of 27,665/15,687 and 2,543,855/2,867,914 positive and negative interactions, respectively. The final protein complex training/test sets consisted of 406/264 complexes. The complete lists of training/test interactions and complexes are available at the supporting web site (http://proteincomplexes.org).

### Calculating protein interaction features from the mass spectrometry datasets

We collected published features from three datasets, Wan *et al.* (Wan et al. 2015), BioPlex (Huttlin et al. 2015) and Hein *et al.* (Hein et al. 2015). Wan fractionation features included four measures of co-fractionation as well as 19 lines of evidence from HumanNet (Lee et al. 2011) and two additional AP-MS datasets (Guruharsha et al. 2011; Malovannaya et al. 2011). Specifically, the co-fractionation measures, as described previously (Havugimana et al. 2012; Wan et al. 2015), included a Poisson noise Pearson correlation coefficient, a weighted cross-correlation, a co-apex score and a MS1 ion intensity distance metric. Each co-fractionation measure was applied to each fractionation experiment, totaling 220 features.

Additional features were taken from Wan *et al.* (Wan et al. 2015). In summary, HumanNet features were originally downloaded from http://www.functionalnet.org/humannet/download.html (file: HumanNet.v1.join.txt). We excluded HS-LC (human literature curated) and HS-CC (human co-citation) evidence codes to remove circularity in the training process. The additional AP-MS fly feature, HGSCore value, was downloaded from Supplemental Table 3 in Guruharsha *et al.* (Guruharsha et al. 2011). The additional AP-MS human feature was based on the MEMOs (core modules) certainty assignments “approved,” “provisional,” and “temporary” downloaded from Supplemental File 1 in Malovannaya *et al*. (Malovannaya et al. 2011), assigning the scores 10, 3 and 1, respectively.

BioPlex AP-MS features were downloaded from: http://wren.hms.harvard.edu/bioplex/data/cdf/150408_CDF_STAR_GRAPH_Ver2594.cdf Specifically, we used the following nine features: NWD Score, Z Score, Plate Z Score, Entropy, Unique Peptide Bins, Ratio, Total PSMs, Ratio Total PSM’s and Unique: Total Peptide Ratio. For the Hein AP-MS data, the features prey.bait.correlation, valid.values, log10.prey.bait.ratio, and log10.prey.bait.expression.ratio were taken from Supplemental Table S2 in Hein *et al*. (Hein et al. 2015).

We generated two additional features for both the BioPlex and Hein AP-MS datasets based on a matrix model interpretation, specifically, the number of experiments a pair of proteins is observed together (pair_count) as well as a *p*-value of two proteins being observed together at random across all AP-MS experiments, as calculated using the hypergeometric distribution as previously described (Hart, Lee, and Marcotte 2007).

### Accurate learning of pairwise protein interactions

Given this feature matrix, we next proceeded to train a support vector machine (SVM) protein interaction classifier. We scaled the feature values using LIBSVM’s (Chang and Lin 2011) `svm-scale` to avoid features with larger numeric range from dominating the classifier. We performed a parameter sweep of the SVM C and gamma parameters using LIBSVM’s cross-validation `grid.py` utility. Training and prediction were calculated using LIBSVM’s `svm-train` and `svm-predict` tools with the ‘probability estimates’ option set to true. Finally, we applied the SVM classifier to all pairs of proteins for which we had data, thereby generating a protein interaction network in which edge weights between protein nodes were set to the SVM’s probability estimate for interacting. We repeated this procedure for combinations of features including only features for individual publications, as well as Wan + BioPlex, Wan + BioPlex + Matrix_Model, Wan + BioPlex + Matrix_Model + Hein and Wan + BioPlex + Matrix_Model + Hein + Matrix_Model (fully integrated). To calculate precision recall curves, we used the python scikit-learn machine learning package (Buitinck et al. 2013).

### Identifying protein complexes by clustering the interaction network

We applied a two stage clustering approach to the protein interaction network to identify clusters of densely interacting proteins, representing our best estimates of protein complexes. First, we sorted the edges of the protein interaction network by their interaction probabilities and selected the top *f* percent of edges, where *f* is a parameter in the range of [0.008, 0.01, 0.015, 0.02, 0.025, 0.03, 0.05] determined by a parameter sweep described below. We applied the ClusterOne algorithm (Nepusz, Yu, and Paccanaro 2012) to the resulting interaction network, specifying minimum size parameter = 2, seed method parameter = ‘nodes’, density in the range [0.2, 0.25, 0.3, 0.35, 0.4] and overlap in the range [0.6, 0.7, 0.8]. For each cluster produced by the ClusterOne algorithm, we refined the clustering by performing a second round of clustering using the MCL algorithm (Enright, Van Dongen, and Ouzounis 2002), specifying the MCL parameter inflation (-I) to be in the range [1.2, 2, 3, 4, 5, 7, 9, 11, 15]. In parallel, we refined each ClusterOne cluster using an alternate second-stage clustering algorithm, the Newman method (Newman 2004). Finally, we removed any protein from the resulting clusters that did not have an edge weight to the remaining proteins in the cluster scoring above the filter parameter, *f*, which occasionally, although rarely, arose through the action of the MCL algorithm.

To objectively optimize the choice of clustering parameters, we performed the two stage clustering process for each combination of parameters, varying *f*, density, overlap and inflation, and selected the cluster set that maximized the F-Grand *k*-clique measure compared to the training set of literature curated complexes. The best-scoring parameters for ClusterOne + MCL were size: 2, density: 0.2, overlap: 0.7, seed_method: nodes, inflation: 7, and *f*: 0.03. The final parameters for ClusterOne + Newman were size: 2, density: 0.4, overlap: 0.7, seed_method: nodes, and *f*: 0.02. Edges that passed the *f* filter corresponding to an interaction probability of 0.26509, were considered high confidence. Finally, we combined the best-scoring two-stage clustering sets (*i.e*., the union of the best performing ClusterOne + MCL and ClusterOne + Newman sets) to form the final estimate of protein complexes.

### Measuring accuracy of the protein complex map by the k-cliques method

We developed a new class of similarity metrics called *k*-cliques for comparing sets of predicted complexes to gold standard complexes. In this method, we consider all observed cliques within a set of clusters. We evaluate precision as whether each clique in our predicted complex set is present or absent in the cliques of the gold standard complex set. Similarly, we evaluate recall according to the presence or absence of cliques from the gold standard complex set in the predicted complex set.

In detail, let *C* be a set of predicted complexes *{c_1_, c_2_,…, c_n_}* and *D* be a set of gold standard complexes *{d_1_, d_2_,…, d_m_}*, where c_i_ and d_j_ are an individual predicted complex and gold standard complex respectively. Let *Q_D_* be the set of protein identifiers in *D* (equation 1).

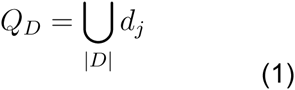

*𝒫* represents the powerset (set of all subsets) and *𝒫_k_* represents the powerset of a given size (eg. *k=*2, all pairwise combinations; *k=3*, all triplet combinations; etc). *A_k_* (equation 2) represents the set of all size *k* cliques in the predicted clusters, *c.*An additional condition on *A_k_* is that the individual cliques overlap with proteins in the gold standard set (*Q_D_*, equation 1), so we only evaluate on proteins that have known complex memberships. The rationale for this so we do not penalize novel predicted complexes as false positives. Similarly, *B_k_*, (equation 3), represents the set of all size *k* cliques in the gold standard complexes set *D*. Note, there is no condition on *B_k_* in terms of protein membership as was done with *A_k_*. This results in an absolute recall measure and evaluates on all complexes in all the gold standard regardless of whether or not there is ample data for those proteins.

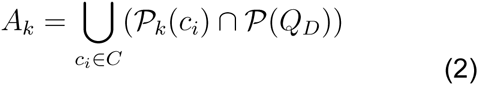

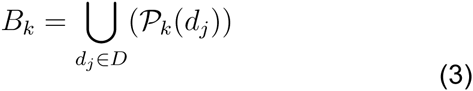

Definitions of *A_k_* and *B_k_* now provide us with a way to compare size *k* cliques in predicted clusters to size *k* cliques in gold standard complexes in a precesion/recall framework. Equations 4,5 and 6 describe the operations of determining true positives (*TP_k_*), false positives (*FP_k_*) and false negatives (*FN_k_*), respectively, for a given clique size *k*.

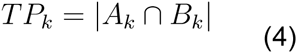

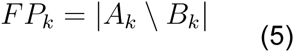

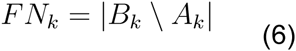

Equations 7 and 8 define precision (*P_k_*) and recall (*P_k_*), and equation 9 defines F-measure (*F_k_*) as the harmonic mean of *P_k_* and *R_k_*

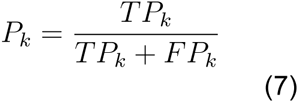

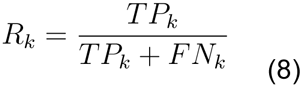

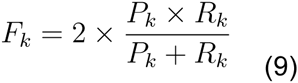

Finally, we define a global F-measure (*F*-*Grand*, equation 10) as the mean of *F_k_*’s, iterating over clique sizes of *k* from 2 to *k* where *k* is the max cluster size of the predicted clustering set *C*.

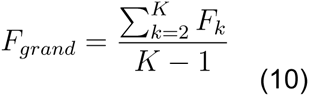

Additionally, we define an alternative global measure that defines weights for each *P_k_* and *R_k_* by the number of clusters, *W_k_*, with size ≥ *k*. This allows for the mitigation of potential bias created by large clique sizes only having a few contributing clusters.

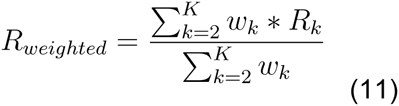

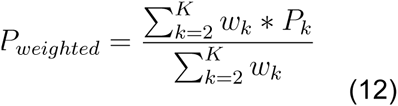

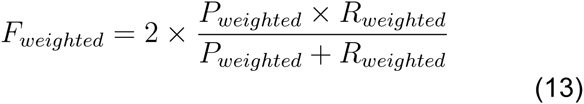

In practice, the sizes of the clique sets (equations 2 and 3) are quite large and computationally intractable to calculate. We therefore randomly sample 10,000 cliques from *A_k_*and *B_k_*when evaluating true positive, false positive and false negative values (equations 4, 5 and 6). Additionally, we add a pseudo-count of 0.00001 to true positive, false positive and false negative values when calculating precision and recall (equations 7 and 8). We have implemented a script to calculate the F-weighted *k*-clique score that is available in our project GitHub repository. An example command line is as follows:

~~~
python    complex_comparison.py  --cluster_predictions    hu_MAP.txt  --
gold_standard testComplexes.txt
~~~

### Measuring overlap with independent protein interaction datasets

In order to assess agreement between our complex map and other protein interaction datasets, we compared the observed overlap of protein interactions to the overlap expected by chance. We chose three datasets that were not integrated into our protein complex discovery pipeline, the yeast two-hybrid dataset from Rolland *et al*. (Rolland et al. 2014), released but unpublished interactions (06/12/2015) from the BioPlex project (“BioPlex” 2016), and an inter-protein crosslinking (1% FDR) dataset from Liu *et al.* (Liu et al. 2015).

For the unpublished BioPlex dataset, we removed all interactions that overlapped with the original published BioPlex interaction set to ensure a disjoint set with our training dataset. For the Liu *et al.* crosslinking dataset, we considered a non-redundant subset by collapsing all inter-protein crosslink interactions for each pair of proteins to just a single interaction.

For each interaction dataset, we generated 1,000 random interaction sets by randomly selecting *M* pairs of proteins where *M* is the number of interactions in that dataset. We then compared the overlap of interactions from our complex map with the random interaction sets to determine a random distribution and calculated a z-score for the overlap of our complex map and the original interaction dataset relative to the random distribution.

### Synaptic bouton complex expression analysis

We downloaded human normalized microarray datasets H0351.2001, H0351.2002, H0351.1009, H0351.1012, H0351.1015, H0351.1016 from the Allen Brain Map (Hawrylycz et al. 2012) [downloaded from: http://human.brain-map.org/static/download]. For each gene in the synaptic bouton complex, we averaged expression values across corresponding probes and calculated Pearson correlation coefficients for each pair of genes. For comparison to a random background distribution, we randomly selected 60 probes from the microarray datasets and calculated Pearson correlation coefficients between the random probes and the genes in the synaptic bouton complex.

For tissue expression analysis of synaptic bouton complex genes, we used RNA-sequencing data for 32 tissues from the Human Protein Atlas (Uhlén et al. 2015) [downloaded: http://www.proteinatlas.org/download/rna_tissue.csv.zip].

### Calculations of tissue specificity, annotation enrichment, and coverage

For comparing tissue specificity, we used reported RNA Tissue Category assignments from the Human Protein Atlas (Uhlén et al. 2015) [downloaded: http://www.proteinatlas.org/download/proteinatlas.tab.gz]. We mapped proteins to HPA entries with RNA tissue category classifications, considering either the entire human proteome, the union of proteins from protein interaction networks of Wan *et al*., BioPlex and Hein *et al*., or the proteins in our final complex map. In order to calculate enriched annotations for each complex, we applied g:Profiler (Reimand et al. 2016) with a Bonferroni *p*-value correction per each complex and excluded electronic annotations from consideration. We used the complete set of proteins in the final protein interaction network as the statistical background. In order to calculate coverage of diseases, we mapped OMIM annotations (Amberger et al. 2015) on to Disease Ontology (Schriml et al. 2012) terms, then selected the top eight disease categories as well as the term “ciliopathies”. We then mapped proteins from our complex map, the Wan *et al.* complex map [downloaded: Supplementary Table 2 from Wan *et al.* (Wan et al. 2015)], the BioPlex complex map [downloaded: Supplemental Table S3 from Huttlin *et al.* (Huttlin et al. 2015)] and Hein *et al.* protein interaction network [downloaded: Supplemental Table S2 from Hein *et al.* (Hein et al. 2015)] onto the Disease Ontology terms. We also evaluated coverage of proteins in the SysCilia Gold Standard Version 1 set of cilia related proteins (downloaded: http://www.syscilia.org/goldstandard.shtml) (van Dam et al. 2013).

### Calculation of prey abundance and network visualization

To calculate percentile ranks of prey abundance for AP-MS raw data, we used the prey abundance measures “zscore” and “prey.bait.correlation” from BioPlex and Hein *et al.* respectively. We ordered each set and calculated the rank percentile using SciPy stats.percentileofscore (Jones et al. 2015) for each pair in the list. Networks of protein complexes were visualized using Cytoscape 3.2.1 (Shannon et al. 2003).

### Morpholinos and mRNA synthesis

Morpholino antisense oligonucleotides (MOs) were purchased from Gene Tools. The *ANKRD55* MO was designed to block splicing using the sequence 5’-TCTGAATCACCTTGAAGCACAAAGA-3’. We used previously validated MOs for *JBTS17*, 5’-TCTTCTTGATCCACTTACTTTTCCC-3’ (Toriyama et al. 2016); and *IFT52*, 5’-AAGCAATCTGTTTGTTGACTCCCAT-3’ (Dammermann et al. 2009). Full length *ANKRD55* cDNA (identified from Xenbase, www.xenbase.org) was amplified from a *Xenopus* cDNA library and subcloned into the vector pCS10 R (derived from pCS107 expression vector) fused with C-terminal GFP. The human *CLUAP1* open reading frame was obtained from the Human ORFeome collection V7.1 and subcloned into the pCS10 R-mCherry vector. Capped mRNAs were synthesized using mMESSAGE mMACHINE (Ambion). mRNAs and MOs were injected into two ventral blastomeres or two dorsal blastomeres at the 4-cell stage to target the epidermis or the neural tissues, respectively. We used each mRNA or MO at the following dosages: *ANKRD55* MO (30 ng for the epidermis and 20 ng for the neural plate), *JBTS17* MO (20 ng), *IFT52* MO (40 ng), *ANKRD55*-GFP mRNA (75 pg), *ANKRD55* mRNA (350 pg for neural tube closure rescue experiment), membrane RFP mRNA (50 pg), and mCherry-*CLUAP1* (100 pg).

### Imaging and analysis

For high-speed live imaging, *Xenopus* embryos injected with *ANKRD55*-GFP and mCherry-*CLUAP1* mRNA were anaesthetized with 0.005% benzocaine at stage 26. High-speed *in vivo* imaging was acquired on a Nikon Eclipse Ti confocal microscope with a 63×/1.4 oil immersion objective at 0.267 sec per frame. Kymographs were calculated using Fiji (Schindelin et al. 2012). Confocal images were collected with an LSM700 inverted confocal microscope (Carl Zeiss) with a Plan-APOCHROMAT 63×/1.4 oil immersion objective. Bright field images were collected using a Zeiss Axio Zoom V16 stereo microscope with Carl Zeiss Axiocam HRc color microscope camera. Neural tube closure quantification was performed using Fiji.

### Plasmids

*CCDC138, CCDC61, TBC1D31, FOPNL, MIB1, PIBF1*, and *SSX2IP* entry ORF clones were obtained from the DNASU Plasmid Repository (Seiler et al. 2014; Grant et al. 2015). *Xenopus laevis* cDNA was prepared by reverse transcription (SuperScriptIII First strand synthesis, Invitrogen), and *KIAA1328, WDR90, TTLL5* cDNAs were PCR-amplified from the library using the following primers:

WDR90 F   caccATGGCTGGAGTCTGGCAG
WDR90 R   TGAATTCTGAATGTCCCACAC
TTLL5 F   caccATGCCCGAAATGTTGCC
TTLL5 R   TTTTCTTTGCCCTTTACTGTCGA
KIAA1328 F   caccATGGATTTACAGAGGCAGCAAG
KIAA1328 R   ACAAATGAAGAAGATCTCCTCTAACATC

PCR products were sub-cloned into Gateway ENTRY clones (pENTR/D-TOPO Cloning Kit, Life Technologies). Destination vectors were modified from destination vector Pcsegfpdest (a gift from the Lawson laboratory) by inserting the **α**-tubulin promoter between the SalI and BamHI sites. Fluorescence protein-tagged expression plasmids were constructed using the LR reaction on entry clones and destination vectors with the Gateway LR Clonase II Enzyme mix (Life Technologies). Expression plasmids (40pg) with centrin-BFP mRNA (100pg) were co-injected into the ventral blastomeres of *Xenopus* embryos at the 4-cell stage and imaged at stage 27.

### Xenopus embryos

*Xenopus* embryo manipulations and injections were carried out using standard protocols. All experiments were performed following animal ethics guidelines of the University of Texas at Austin, protocol number AUP-2015–00160.

## Acknowledgements

This work was supported by grants from the NIH (F32 GM112495 to K.D.; 1R01 HL117164 to J.B.W.; R21 GM119021, R01 HD085901 to J.B.W. and E.M.M., and DP1 GM106408, R01 DK110520 to E.M.M.), NSF and CPRIT (to E.M.M.), and the Welch foundation (F-1515, to E.M.M.). Protein interactions and complexes have been deposited in the BioGRID database (Stark 2006) (accession id in progress) and can be searched interactively or downloaded at http://proteincomplexes.org Supporting computer code for the full protein interaction mapping and analysis pipeline is available at: https://github.com/marcottelab/protein_complex_maps_public

## Author Contributions

K.D. and E.M.M designed project. K.D. developed code and performed data analysis. C.L, R.L.H., F.T., Y.M and J.B.W designed and performed validation experiments. B.B. contributed code. C.D.M. performed analysis. K.D., J.B.W and E.M.M drafted manuscript. All authors discussed results and contributed edits.

## Expanded View

**Supplemental Figure 1**: hu.MAP outperforms published complex maps on leave-out set of gold standard complexes

**Supplemental Figure 2**: hu.MAP recapitulates IFT A and IFT B complexes

**Supplemental Figure 3**: ANKRD55 localization and movement along axonemes is consistent with being a component of the IFT-B particle

**Supplemental Table 1**: Protein and interaction overlap for published networks

**Supplemental Table 2**: hu.MAP complexes

**Supplemental Table 3**: hu.MAP interaction network

**Supplemental Table 4**: hu.MAP complex annotation enrichments

**Supplemental Table 5**: hu.MAP ciliary complexes

**Supplemental Movie 1**: ANKRD55 traffics along the axoneme of X. *laevis* cilia

**Supplemental Movie 2**: ANKRD55 co-migrates along axonemes with known IFT-B component CLUAP1

**Supplemental Movie 3**: Single axoneme view of ANKRD55 co-migrating with known IFT-B component CLUAP1

## Figure Legends

**Supplemental Figure 1:**
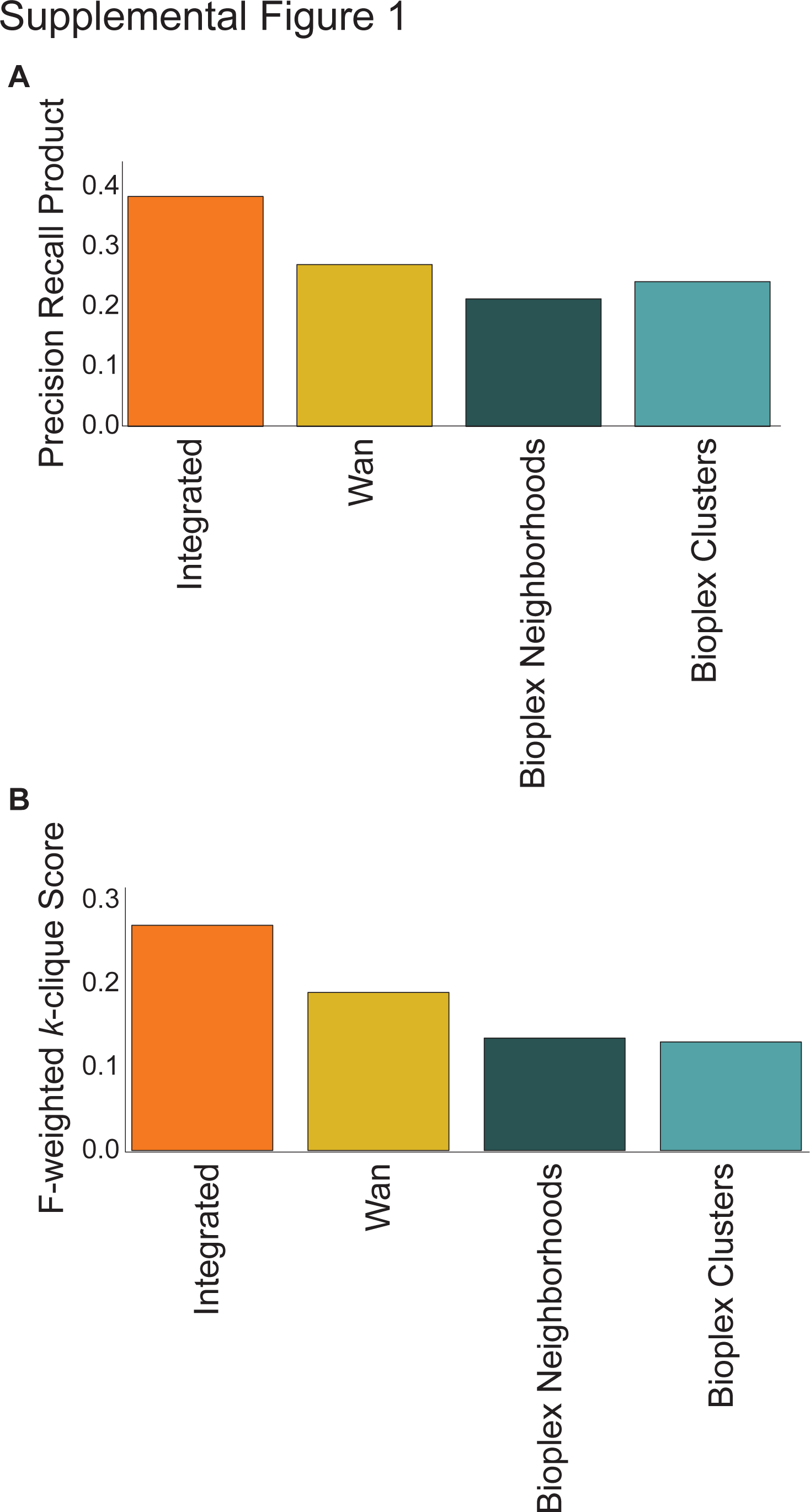
hu.MAP outperforms published complex maps on leave out set of gold standard complexes. A. Comparison of hu.MAP and published complex maps to leave out set using precision recall product measure (Song and Singh 2009). B. Comparison of hu.MAP and published complex maps to leave out set using F-weighted *k*-clique score.

**Supplemental Figure 2:**
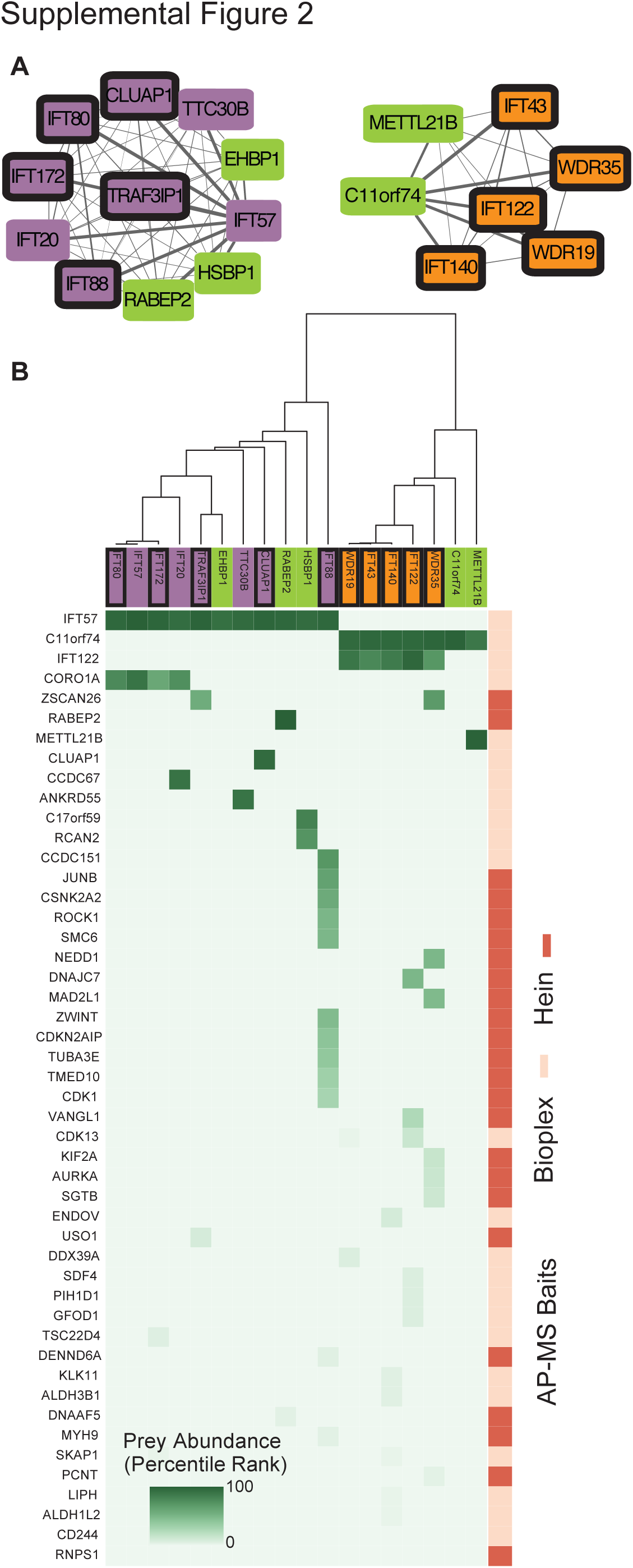
hu.MAP recapitulates IFT A and IFT B complexes. A. Network view of IFT A and IFT B complexes. Node colors follows **Figure 5** conventions. B. Matrix of AP-MS experiments show IFT A and IFT B are well separated and supported by multiple experiments.

**Supplemental Figure 3:**
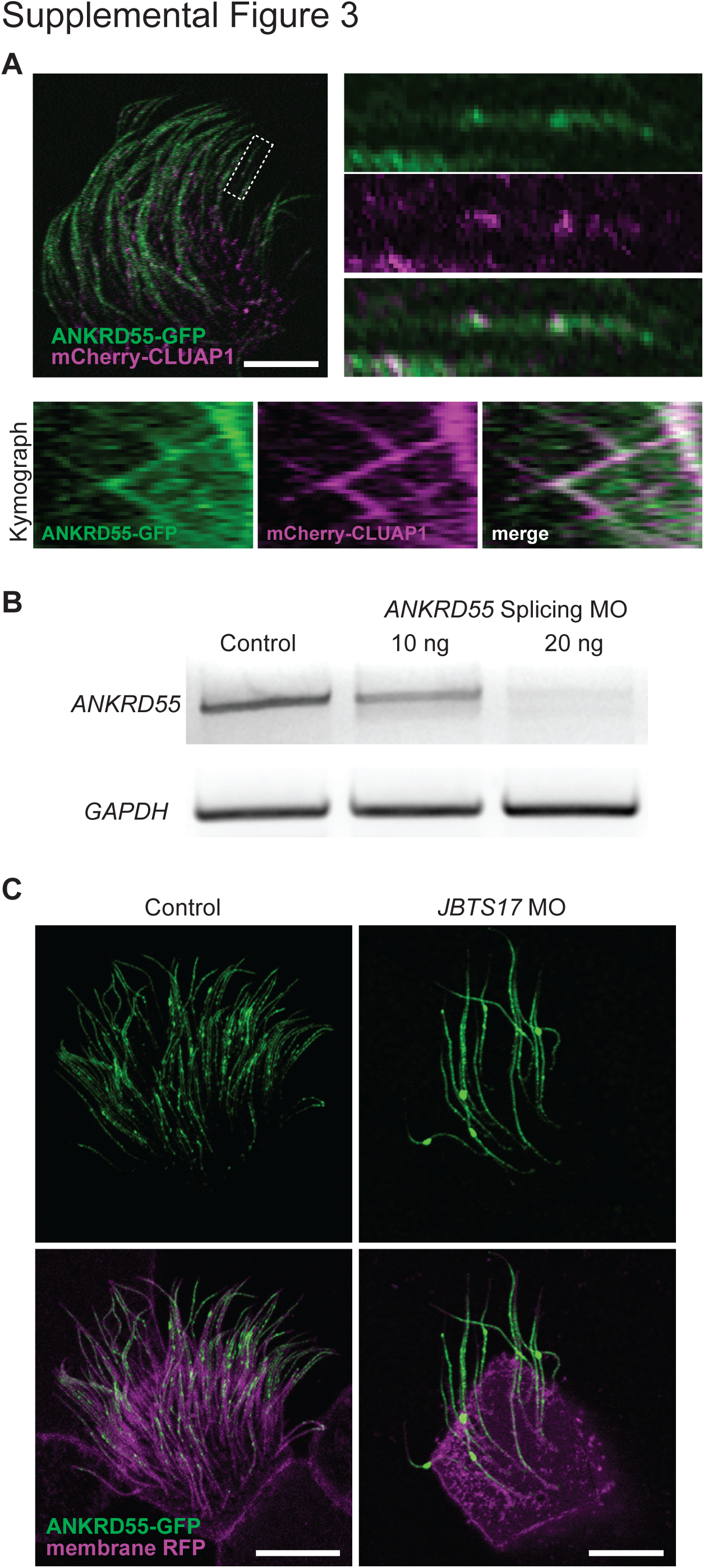
ANKRD55 localization and movement along axonemes is consistent with being a component of the IFT-B particle. A. Two color kymograph generated by co-expression of ANKRD55-GFP (green) and mCherry-CLUAP1 (magenta) reveals that ANKRD55 travels along axonemes in association with other IFT proteins. Scale bar: 10 um. B. RT-PCR demonstrates the efficiency of *ANKRD55* MO to disrupt splicing of *ANKRD55* mRNA in *Xenopus* embryos. GAPDH is used as control. C. Morpholino knockdown of *JBTS17*, known to specifically effect IFT-B localization, results in the accumulation of ANKRD55-GFP in axonemes (Green: ANKRD55-GFP, Magenta: membrane RFP). Scale bar: 10 um.

**Supplemental Movie 1: ANKRD55 traffics along the axoneme of X. *laevis* cilia (green, ANKRD55-GFP; magenta, membrane RFP).**

**Supplemental Movie 2: ANKRD55 co-migrates along axonemes with known IFT-B component CLUAP1 (green, ANKRD55-GFP; magenta, CLUAP1-RFP).**

**Supplemental Movie 3: Single axoneme view of ANKRD55 co-migrating with known IFT-B component CLUAP1 (green, ANKRD55-GFP; magenta, CLUAP1-RFP). Left = proximal, right = distal.**

